# Pericyte Bridges in Homeostasis and Hyperglycemia: Reconsidering Pericyte Dropout and Microvascular Structures

**DOI:** 10.1101/704007

**Authors:** Bruce A. Corliss, H. Clifton Ray, Richard Doty, Corbin Mathews, Natasha Sheybani, Kathleen Fitzgerald, Remi Prince, Molly Kelly-Goss, Walter L. Murfee, John Chappell, Gary Owens, Paul Yates, Shayn M. Peirce

**Author notes:** **Funding:** NIH R21 EY028868-01, NIH-U01AR069393, NIH-U01HL127654, The Hartwell Foundation, The Stanford Allen Discovery Center (to S.M.P.). **Corresponding Author:** Shayn M. Peirce, Ph.D., Professor, Department of Biomedical Engineering, 415 Lane Road, University of Virginia, Charlottesville, VA 22908.

## Abstract

Diabetic retinopathy threatens the vision of a third of diabetic patients. Progression of the disease is attributed to the dropout of pericytes, a cell type that enwraps and stabilizes the microvasculature. In tandem with this presumptive pericyte dropout, there is enriched formation of structures assumed to be remnants of collapsed or regressed vessels, previously classified as acellular capillaries, string vessels, and basement membrane bridges. Instead of endothelial cells, we show that pericytes colocalize with basement membrane bridges, and both bridging structures are enriched by cell-specific knockout of KLF4 and reversibly enriched with elevation of Ang-2, PDGF-BB, and blood sugar. Our data suggests that pericyte counts from retinal digests have misclassified pericyte bridges as endothelial structures and have exaggerated the role of pericyte loss in DR progression. In vivo imaging of corneal limbal vessels demonstrates pericyte migration off-vessel, implicating pericyte movement in formation of pericyte bridges and pathogenesis of diabetic retinopathy.

## Introduction

In diabetes, the malfunction of pericytes, cells that enwrap capillaries and considered to be key effectors in microvascular remodeling^1^, is thought to contribute to endothelial dysfunction and microvessel dropout across tissues including kidney^2^, skeletal muscle^3^, liver^2^, brain^2^, heart^4^, and retina^2^. Diabetic retinopathy is the primary condition responsible for working age blindness in the developed world^5^. Its associated pathologies, including aberrant vessel remodeling, pathological angiogenesis, progressive fibrosis, and retinal detachment^5^, have flawed treatment options. Laser photocoagulation treatment sacrifices peripheral and night vision for temporarily slowing pathological angiogenesis^6^. Anti-VEGF treatment can require monthly ocular injections resulting in mixed clinical outcomes: including lack of clinically significant visual improvement^7^, poor patience compliance^7^, and induction of retinal ganglion and neuronal cell death^8^. Better methods are needed for modulating microvascular cell behaviors to improve clinical outcomes for diabetic retinopathy patients.

Pericytes are considered an effector cell for microvascular remodeling and enwrap capillaries, maintaining close physical contact via cell soma and extended cellular processes within the vascular basement membrane^1^. However, some studies have observed pericyte-like cells bridging across two or more adjacent capillaries in both homeostatic^9^ and pathological conditions such as diabetes, where their frequency is elevated^10^. These studies have framed the existence of pericyte-like bridges as evidence of pericyte detachment, where it is assumed that a fully-attached pericyte migrates (or begins to migrate) away from the capillary on which it resides and extends cell processes or its entire cell soma to form a bridge from one capillary to another^10–13^. Previous research has indicated that these bridging cells can colocalize with basement membrane structures that also span across, or bridge, adjacent capillaries^9,14^. These stand-alone (i.e. cell-free) basement membrane structures have been referred to in the literature as acellular capillaries^15^, intervascular bridges^9^, basal lamina and collagen-IV (Col-IV) sleeves^16^, and string vessels^15^. They also appear more frequently in pathological settings than in homeostasis, and some have presumed these basement membrane bridges to be residual structures left by collapsed and regressed capillaries (for review, please see^15^). Together, these observations and assertions point to unanswered questions about the origin, functional significance, longevity, and reversibility of these cellular and acellular cross-capillary bridges, including whether they are formed by migrating pericytes that deposit basement membrane or from remnants of regressed vessels^9^; and, given their increased prevalence in diabetic conditions, whether their frequencies are reversibly tunable by exogenous interventions, and, if so, on what time frame.

To begin to answer these questions, we first evaluated the phenotypic identity of these cell bridges using immunolabeling of a pericyte-specific marker and confirmed with a lineage-tracing genetic reporter mouse driven by the same promoter, which enabled pericytes and their progeny to be fluorescently tagged with a high degree of specificity^17^. Given that all prior studies of these bridging cells have relied exclusively on non-specific proteins like NG2 and PDGFRβ^18^ (which can be expressed by other cell types^19^, such as macrophages and microglia), we regarded this as a critical first step. Next, we tested the hypothesis that pericyte bridges in the retina constitute a subset of pericytes that can be reversibly enriched with multiple interventions that have relevance to diabetic retinopathy and pericyte dynamics, including: pathological elevation of blood glucose, delivery of exogenous pericyte chemokines that are upregulated in diabetes^20,21^ and targeted in clinical trials^22,23^, and pericyte specific knockout of a gene that has been associated with cell migration^24^. Since the establishment and maintenance of basement membrane is an integral role for pericytes^25^, we also sought to examine the frequency of basement membrane bridges and their colocalization with pericyte bridges. Characterizing colocalization of basement membrane bridges with pericytes and endothelial cells could give insight to their origin and function within the microvasculature. The existence of pericyte bridges in healthy conditions suggest an unknown and potentially important role for them in the maintenance of homeostasis, while the enriched density of pericyte bridges in response to hyperglycemia could represent a putative reversible vascular remodeling event in diabetes that could have protective effects if appropriately modulated.

## Methods

### Mouse Strains and Protocol

All procedures were approved by the Institutional Animal Care and Use Committee at the University of Virginia, and completed in accordance with our approved protocol under these guidelines and regulations. With the lineage tracing and live imaging, *Myh11*-CreER^T2^ ROSA floxed STOP tdTomato mice used, bred from *Myh11*-CreEr^T2^ Mice (Jackson Laboratory, Cat. # 019079) and ROSA floxed STOP tdTomato mice (Jackson Laboratory, 007914), all on C57bl/6J background. Lineage specific knockout of KLF4 was investigated with *Myh11*-CreER^T2^ ROSA floxed STOP eYFP *Klf4*^fl/lf^ with *Myh11*-CreER^T2^ ROSA floxed STOP eYFP *Klf4*^WT/WT^ used as control graciously provided by the Gary Owens Lab (University of Virginia, Charlottesville) and were treated with tamoxifen as done previously^26^ (Supplementary Materials 1). All other mice used were C57Bl6J (JAX stock #000664, Bar Harbor, ME).

STZ treatment was conducted as done previously^27^ (Supplementary Materials 1). Mice injected intravitreally with Ang2 or PDGF-BB (Supplementary Materials 2) and were examined at day 4 and additionally day 28: long after exogenous protein had dissipated based on the short half-lives of Ang2^28^ and PDGF-BB^29^. Mice were sacrificed and immunostained using previously developed techniques^30,31^ (See Supplementary Materials 3 and Supplementary Tables 1-2 for antibodies used).

### Quantifying Microvascular Structure and Pericyte Phenotype

Cell counts of pericyte association state with the vasculature^32^ was quantified using Fiji’s Cell Counter plugin^33^ in a blinded fashion. Vessel structure was analyzed with software written in MATLAB using previously developed methods^34^ (Supplementary Materials 4).

### Data Acquisition, Statistics, and Sampling

Two-tailed tests were used with significance level was set to *α* = 0.05. For the plots, ‘*’ denotes p<0.05, ‘**’ denotes p<0.01, and ‘***’denotes p<0.001. See Supplementary Materials 5 for further details.

## Results

### Pericyte Marker NG2 Colocalizes with Basement Membrane Bridges in Homeostasis

Previous research shows that basement membrane bridges are found in healthy homeostatic conditions in the retina^9^, suggesting some form of ongoing remodeling of the basement membrane. We hypothesize that in the homeostatic retina, endothelial cells are not associated with these structures, while pericytes are. Immunostaining of retinal digests, an assay used to isolate the vasculature via enzymatic digestion where only endothelial cells, pericytes, and basement membrane bridges remain^10^, reveals that these thin basement membrane structures are Col-IV+, but negative for pan-endothelial marker CD31^35^ and retina-specific endothelial marker CD105^36^ (Fig. 1A-D), confirming previous findings^35^. In wholemount immunostained retinas, these Col-IV+ bridges are also colabeled with other known markers of the basement membrane, including fibronectin, laminin, and IB4 lectin (Fig. 1E-G). These basement membrane bridges, and the associated cell soma often found connected to them, appear to morphologically match the structures classified as acellular capillaries in non-specific cell staining of retinal digests (Fig. 1H-I). Basement membrane bridges are also observed in retinas of humans, macaque, and rabbit (Supplemental Fig. 1A-D).

**Figure 1:**
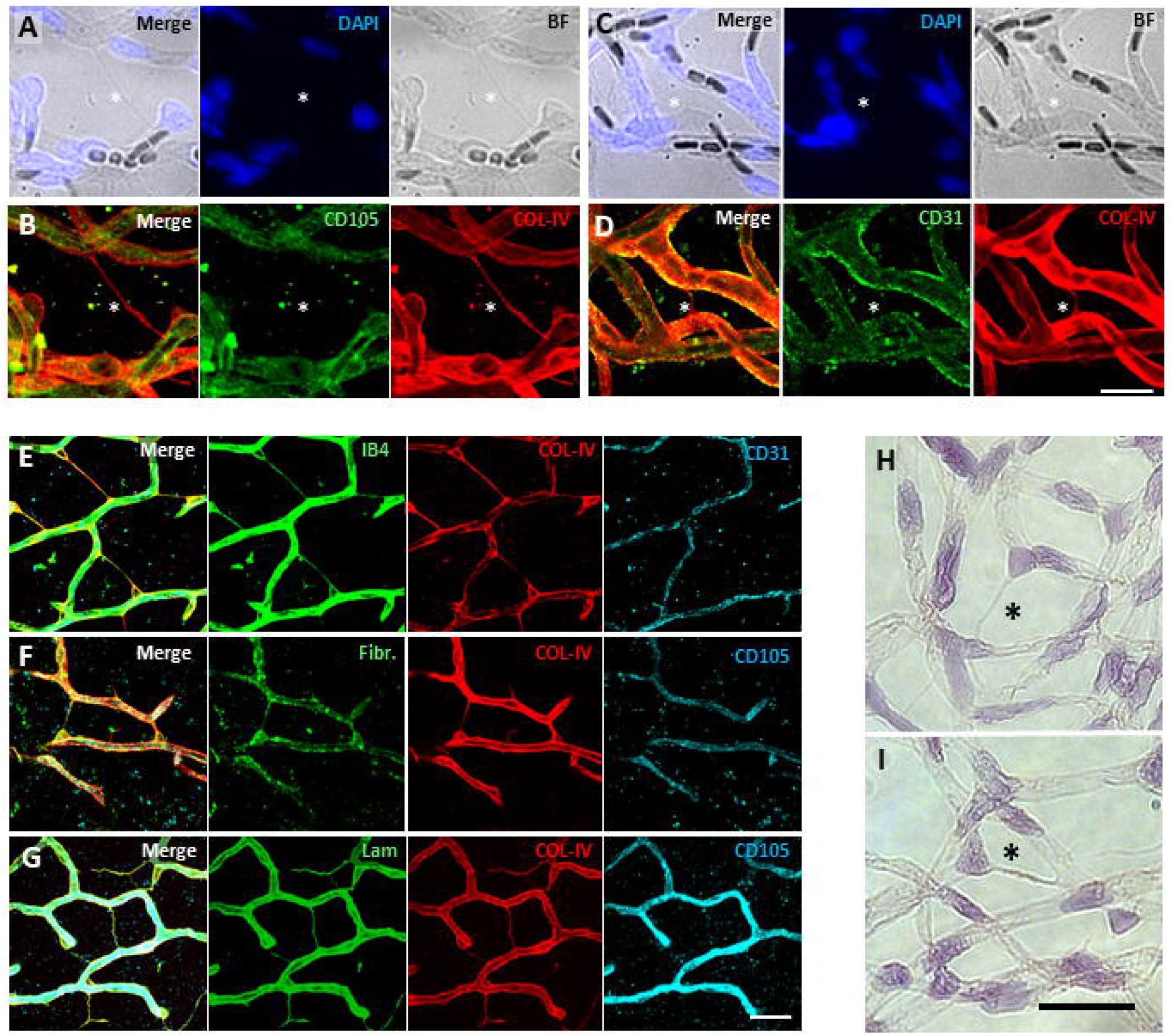
Acellular capillaries in murine retina colabel with basement membrane markers but lack endothelial markers. (**A**) Brightfield image of acellular capillary (star) between two fully formed capillaries with (**B**) field of view imaged fluorescently with anti-CD105 (green) and anti-COL-IV (red). (**C**) Brightfield of acellular capillary (star) with (**D**) same location fluorescently imaged with anti-CD31 (green) and anti-COL-IV (red) (scale bar 15 μm). (**E**) Fluorescent confocal image of wholemount retina in deep plexus stained with isolectin-IB4 (green), anti-collagen-IV (red) and anti-CD31 (cyan). (**F**) Retinal deep plexus labeled with isolectin-IB4 (green), anti-fibronectin (red) and anti-CD105 (cyan). (**G**) Retinal deep plexus labeled with isolectin-IB4 (green), anti-laminin (red) and anti-CD105 (cyan) (scale bar 25 μm). (**H, I**) Transmission light image of retinal digest stained with hematoxylin and eosin, with structures previously referred to as acellular capillaries (star) (scale bar 15 μm).

We examine healthy murine tissue and characterize these structures in the deep retinal plexus. Representative images of immunostaining of homeostatic retina (Fig. 2A) reveals capillaries and off-vessel bridges labeled with Col-IV+ basement membrane, as well as no signs of vessel formation or regression, as expected in homeostasis. When the diameter of basement membrane segments (blood vessels and basement membrane bridges) of the microvascular network was grouped based on marker expression, endothelial colabeled segments (CD31+ and CD105+) had a diameter consistent with capillaries at 5.64 ± 0.08 μm while non-endothelial segments had a 60.5% reduced diameter of 2.23 ± 0.08 μm (Fig. 2B p=2.99E-12). The complete separation of diameter values between endothelial and non-endothelial segments, with a lack of segments that have an intermediate diameter transitioning between the two structures, hints that endothelial cells may not be related with the occurrence of these structures. Furthermore, if the marker expression of Col-IV+ basement membrane bridge segments are examined (diameter less than 3.5 μm), none of the segments colabel with endothelial cell markers, while 20.5 ± 1.4% of them colabel with NG2 (denoting the presence of an active pericyte cell process), and 79.5 ± 1.4% colabel with neither (Fig. 2C, p=3.70E-27).

**Figure 2:**
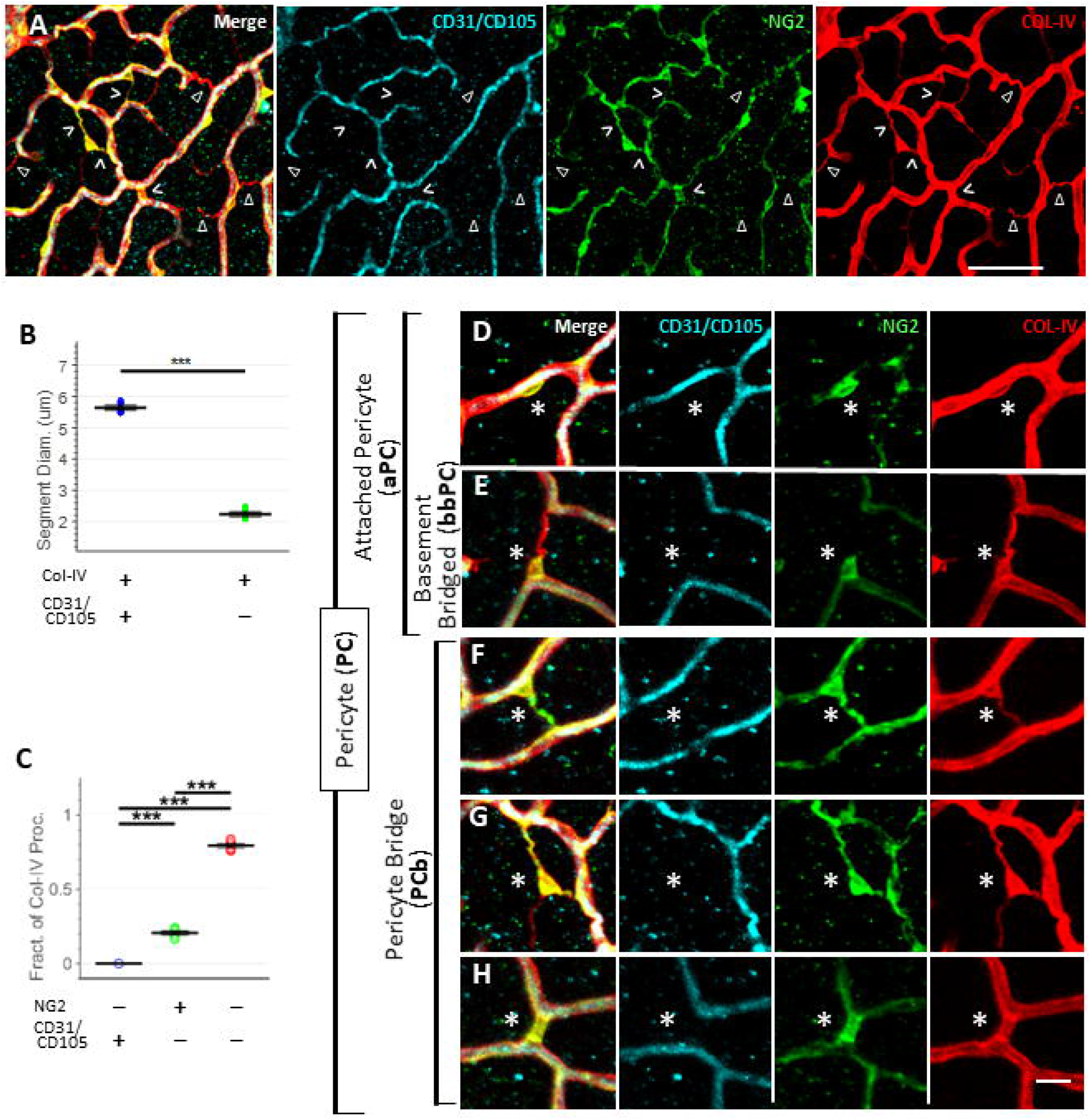
Basement membrane bridges, a subset of which colabel with the pericyte marker NG2, represent distinct morphological structures compared to lumenized vessels. (**A**) Various off-vessel Col-IV tracks (red, closed arrow) are observed bridging CD31/CD105 blood vessels (cyan) in deep plexus of the retina, a subset of which colabel with NG2 (green, open arrow) (scale bar 50 μm). (**B**) Comparison of diameter of collagen-IV segments that colabel with and without the endothelial markers CD31 and CD105 (Welch’s t-test, N=6 mice). (**C**) Fraction of colabeled basement membrane bridges (Col-IV structures less than 3.5 um diameter) that colabel with pericyte marker NG2, endothelial markers CD31/CD105, and neither (1-way Kruskal Wallis with Bonferroni correction, N=6 mice). Classification of various pericyte phenotypes (star) in relation to the vascular network, including (**D, E**) a fully attached pericyte, some of which are connected to empty basement membrane bridges, and (**F-H**) various types of pericyte bridges (scale bar 15 um).

A range of NG2+ cell morphologies were found, which we classify as either an attached pericyte with cell soma and all cell processes associated with a vessel (Fig. 2D, E) or a pericyte bridge with a cell soma or process extending partially off-vessel (Fig. 2F-H). Additionally, we divided attached pericytes into a subgroup with those that are connected by an off-vessel basement membrane bridge (basement membrane bridged pericyte) that lack colabeling with pericyte and endothelial cell markers, giving an impression of an empty basement membrane track (Fig. 2E).

### Pericyte Bridges Express the Smooth Muscle Cell and Pericyte Specific Marker Myh11

Previously, the Myosin Heavy Chain 11 (Myh11) promotor has been used in an inducible lineage tracing reporter mouse model to track the lineage of smooth muscle cells^37^. Myh11 lineage cells also colabel with the majority of NG2 or PDGFRβ expressing pericytes^17^, and have been used to study them. We hypothesized that pericyte bridges are marked in this mouse model, and the exclusivity of Myh11 expression could be leveraged to visualize pericyte morphology with higher confidence of cell identity than with NG2 or PDGFRβ as markers. Post tamoxifen induction in the *Myh11*-CreER^T2^ ROSA floxed STOP tdTomato (“Myh11-RFP”) mouse model in the deep retinal plexus, RFP+ Myh11 lineage (Myh11-Lin(+)) cells colabeled with NG2+ cells that bridged between blood vessels marked with CD31 (Fig. 3A-B), CD34, and CD105 (Supplementary Fig. 1E-F). Since RFP expression denotes Myh11 expression induced during the tamoxifen treatment period ending 4 weeks prior to sacrifice, these Myh11-Lin(+) cells could have lost Myh11 expression during the chase period. However, we find these bridging cells are also marked when immunostained directly for the Myh11 protein (Fig. 3C). The fact that Myh11 expression is found in these bridging cells in tandem with NG2 expression and colocalizing with a Col-IV+ basement membrane argues that pericyte bridges maintain their pericyte identity and Myh11 can serve as a marker that includes this pericyte subpopulation. Leveraging the ability of this mouse model to label pericyte bridges endogenously, we were able to readily visualize pericyte bridges in other tissues via simple wholemount in various tissues (Fig. 3D-K), suggesting that this morphology represents a fundamental pericyte phenotype found across vascularized tissues.

**Figure 3:**
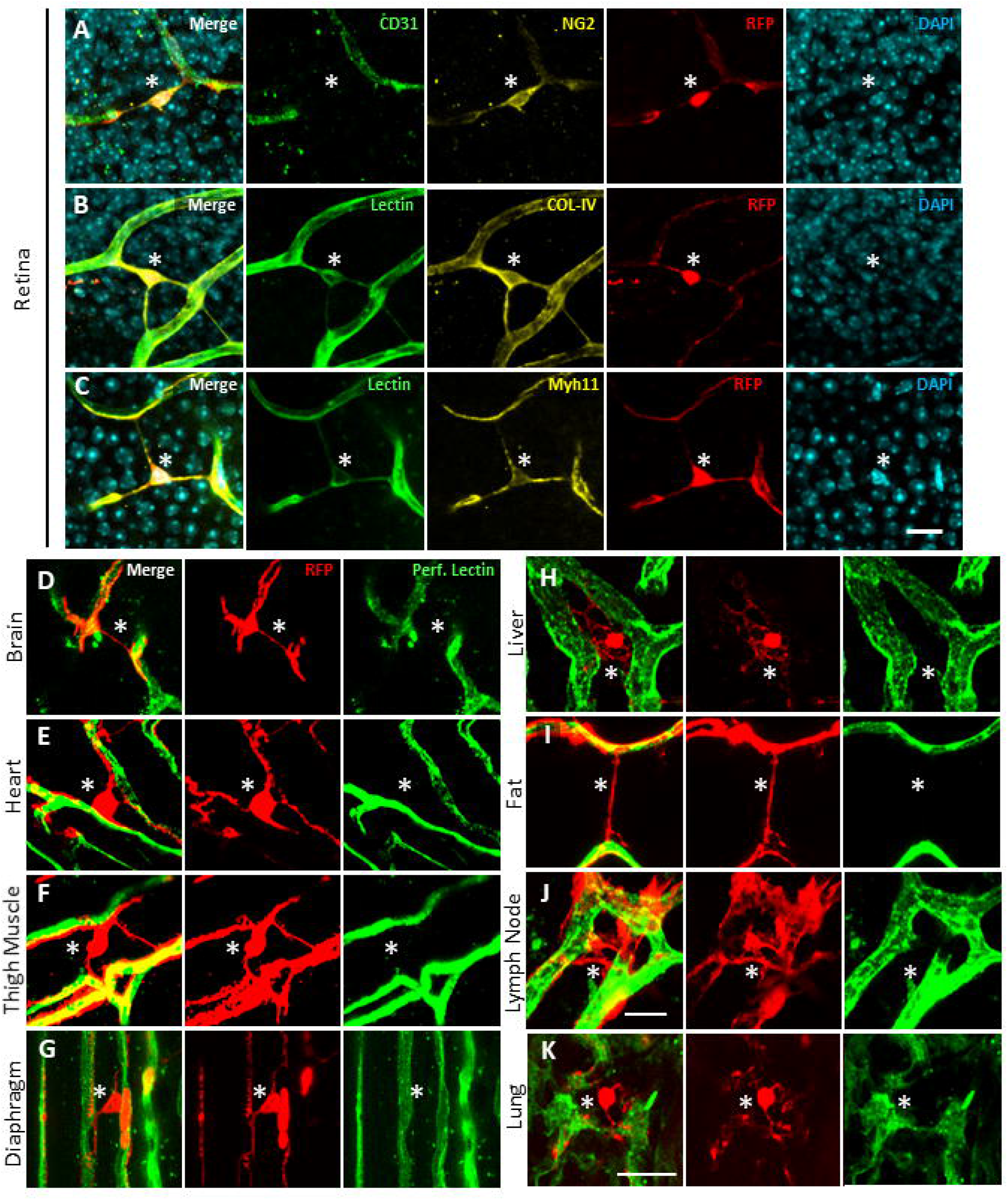
Pericyte bridges express Myh11, are of Myh11 lineage, and found across various tissues in quiescence. (**A**) Pericyte bridge of Myh11-lineage labeled with anti-CD31 (green), anti-NG2 (yellow), RFP (red), and DAPI (cyan). (**B**) Pericyte bridge labeled with IB4 lectin (green), anti-collagen-IV (yellow), RFP (red), and DAPI (cyan). (**C**) Pericyte bridge labeled with IB4 lectin (green), anti-Myh11 (yellow), RFP (red), and DAPI (cyan) (scale bar 15 um). Myh11-lineaged RFP+ cells (red) imaged with perfused IB4 lectin (green) in (**D**) brain, (**E**) heart, (**F**) thigh muscle, (**G**) diaphragm, (**H**) liver, (**I**) inguinal fat, (**J**) lymph node, and (**K**) lung (scale bar 15 um).

### Short-Term and Long-Term Hyperglycemia Elevates While Insulin Treatment Reduces Pericyte Bridge Density in STZ-induced Diabetes

Previous data from retinal digests suggests that basement and pericyte bridges are enriched over the long-term in diabetic conditions: we examined if acute short-term hyperglycemia can modulate bridge density in a reversible fashion by imaging the deep plexus retinal vasculature 1 week and 2 weeks post-streptozotocin (STZ) treatment, STZ treatment with sustained insulin treatment via sub-cutaneous osmotic pump, and vehicle control (Fig. 4A). Mice with STZ-induced diabetes had a 20.9% increase in enriched pericyte bridge density compared to vehicle at day 7 (Fig. 4B) mirrored by a 4.3% reduction in attached pericyte density (Fig. 4C, p=4.27E-3). At day 14, pericyte bridges in diabetic mice were enriched 50.5% and attached pericytes decreased 13.8% (p=4.65E-8). Insulin treatment conferred partial rescue, leading to a 49.3% reduction in pericyte bridge density compared to untreated diabetic mice, along with 8.3% increase in attached pericytes (p=3.90E-5). We observed a similar trend towards enriched basement membrane bridged pericytes with STZ treatment at day 7 (Fig. 4D), and a 34.9% increase in density at day 14 (p=7.21E-5), along with a trend towards 28.2% reduction with insulin treatment compared to untreated diabetic condition (p=6.68E-2). However, there was no change between diabetic and vehicle in the total NG2+ population for days 7 (Fig. 4E, p=0.653) and 14 (p=0.510). Basement membrane bridges marked with Col-IV, along with the subset of those that were colabeled with NG2, displayed the same trends as pericyte bridges with diabetes and insulin treatment (Fig. 4F-G). There was no evidence of angiogenesis or regression, with no changes to vessel length density (Fig. 4H, day 7 p=0.848, day 14 p=0.599), branchpoints per vessel length (Fig. 4I, day 7 p=0.987, day 14 p=0.712), segment tortuosity (Fig. 4J, day 7 p=0.131, day 14 p=0.244), or vessel diameter (Fig. 4K, day 7 p=0.442, day 14 p= 0.423). In support of perivascular remodeling, the density of pericyte bridges, basement membrane bridged pericytes, and off vessel bridges all correlated with mouse blood glucose levels across groups and timepoints at the time of sacrifice (Supplementary Fig. 2D-G). Blood glucose and mouse weight confirms hyperglycemia for each study group (Supplementary Fig. 2A-C); representative images of retinas harvested from hyperglycemia and control mice at days 7 (Supplementary Fig. 2 H, I) and 14 (Fig. 4L-O) provided. Similar patterns are observed with hyperglycemia over the longer term of months with STZ-induced diabetes (Supplementary Fig. 3A-Q, Supplementary Fig. 4A-C, Supplementary Data 1) and with genetic knockout of insulin in the Akita mouse strain^11^ (Supplementary Fig. 5A-Q, Supplementary Data 2).

**Figure 4:**
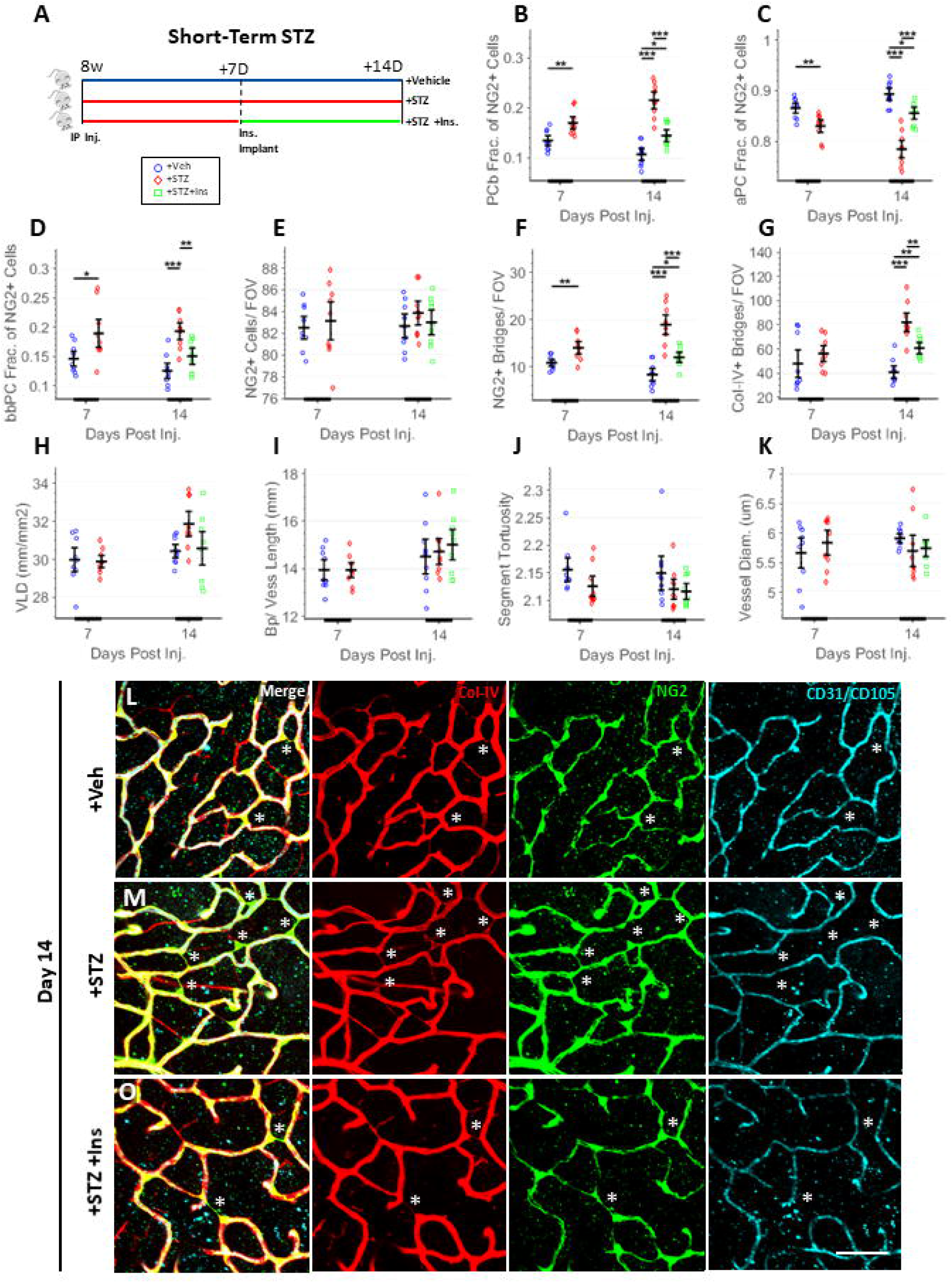
Short term STZ-induced hyperglycemia has enriched pericyte bridge density normalized with short-term insulin treatment over a static vessel network structure. (**A**) Experiment design. In the deep retinal plexus, quantification of pericyte morphology, including (**B**) fraction of NG2+ cells with pericyte bridging phenotype, (**C**) fraction of NG2+ cells with attached pericyte phenotype, (**D**) fraction of NG2+ cells with basement membrane bridged phenotype, (**E**) total NG2+ cells per field of view, (**F**) NG2+ bridges per field of view, and (**G**) all Col-IV+ bridges per field of view. Vessel network morphology quantified with (**H**) vessel length density (mm/mm2), (**I**) branch points per vessel length, and (**J**) vessel segment tortuosity, and (**K**) vessel diameter (day 7: unpaired t-test, day 14: 1-way ANOVA with Tukey multiple comparisons, N=9 mice, FOV 530 μm). (**L-O**) Representative images of retinal deep plexus at day 14 from each treatment group, with anti-Col-IV (red), anti-NG2 (green), along with anti-CD31 and anti-CD105 (cyan), with annotated pericyte bridges (star) (scale bar 50 um).

### Injection of Recombinant PDGF-BB and Ang2 Transiently Elevates Pericyte Bridge Density

Platelet Derived Growth Factor BB (PDGF-BB), a pericyte chemokine elevated in diabetes^38^, binds to PDGFRβ expressed by pericytes, but not endothelial cells^39^. We hypothesized that addition of exogenous PDGF-BB would result in an enriched density of pericyte bridges with no change to total pericyte density. Four days post PDGF-BB injection (Fig. 5A), pericyte bridge density was enriched 47.1% (Fig. 5B) and attached pericytes reduced 19.2% relative to vehicle control (Fig. 5C, p=2.39E-5). At day 28, pericyte bridge density recovered to basal levels compared to control with a trending increase of 11.0%, while attached pericytes were qualitatively reduced 2.2% (p= 0.0866). In contrast with Ang2 stimulus, basement membrane bridged pericyte density was reduced 57.5% at day 4 compared to control (Fig. 5D, p=1.65E-3) and restored by day 28 (p=0.770). Total NG2-labeled cell density remained constant at both days 4 (Fig. 5E, p=0.583) and 28 (p=0.342). Col-IV bridges and the subset colabeled with NG2 followed trends similar to pericyte bridge density across study groups (Fig. 5F-G). No angiogenesis was observed per vessel length density (Fig. 5H, p=0.459), a modest 11.7% increase in branchpoints per vessel length (Fig. 5I, p=0.0147), no change to segment tortuosity (Fig. 5J, p=0.202), and no change to vessel diameter (Fig. 5K, p=0.829). Representative images at days 4 (Fig 5L-M) and 28 (Supplementary Fig. 7A-B) are provided. Similar patterns are observed with injection of Ang2 (Supplementary Fig. 6A-M, Supplementary Fig. 7C-D, Supplementary Data 3).

**Figure 5:**
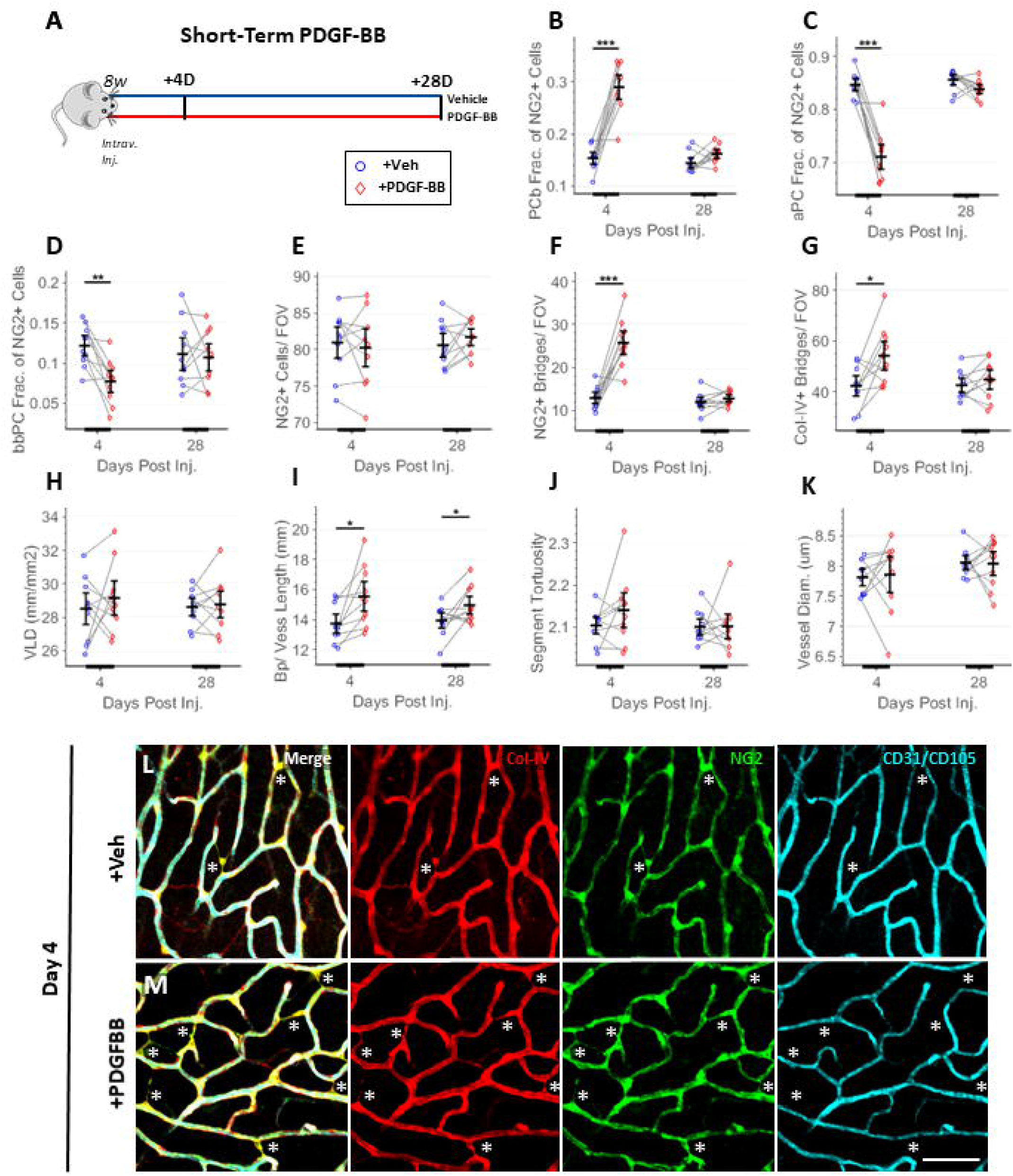
Intravitreal injection of PDGF-BB has transiently enriched pericyte bridge density over a morphologically static vessel network. (**A**) Experiment design. In the deep retinal plexus, quantification of pericyte morphology, including (**B**) fraction of NG2+ cells with pericyte bridge phenotype, (**C**) fraction of NG2+ cells with attached pericyte phenotype, (**D**) fraction of NG2+ cells with basement membrane bridged phenotype, (**E**) total NG2+ cells per field of view, (**F**) NG2+ bridges per field of view, and (**G**) all COL-IV+ bridges per field of view (N=10 mice). Vessel network morphology quantified with (**H**) vessel length density (mm/mm2), (**I**) branch points per vessel length, (**J**) vessel segment tortuosity, and (**K**) vessel diameter (paired t-test at each timepoint, N=10 mice, 530 μm FOV). (**L, M**) Representative images of retinal deep plexus at day 4 from each treatment group, with anti-COL-IV (red), anti-NG2 (green), along with anti-CD31 and anti-CD105 (cyan), with annotated pericyte bridges (star) (scale bar 50 um).

### Knockout of KLF4 Results in Elevated Pericyte Bridge Density

In vascular smooth muscle cells, KLF4 can negatively regulate cell migration^24,40^, and its expression in Myh11 lineaged cells has been shown to alter smooth muscle cell phenotype^26^. To elucidate a possible molecular mechanism that modulates pericyte bridge formation, mice with Myh11 lineage specific inducible KLF4-KO (Myh11-Cre^Ert^EYFP^+/+^KLF4^fl/fl^) were characterized relative to wildtype (WT) (Myh11-Cre^Ert^EYFP^+/+^KLF4^WT/WT^) littermate control mice (Fig. 6A). YFP+ pericyte bridges were enriched by 48.2% in the deep plexus of homeostatic retina in KLF4-KO mice compared to WT (Fig. 6B), while YFP+ attached pericytes were reduced by 8.7% (Fig. 6C, p=1.22E-5). YFP+ basement membrane bridged pericytes were enriched 41.5% within KLF4-KO retinas (Fig. 6D, p=1.61E-3). YFP+ cell density was not altered in the knockout mice (Fig. 6E, p=0.699). Laminin bridges and the subset colabeled with YFP follow similar trends as pericyte bridge density across study groups (Fig. 6F-G). There was no evidence of angiogenesis as measured by vessel length density (Fig. 6H, p=0.451), no change to branchpoints per vessel length (Fig. 6I, p=0.395), enriched vessel segment tortuosity in the knockout (Fig. 6J, p=7.27E-3), and no change to vessel diameter (Fig. 6K, p=0.264). Representative images are shown at 14 weeks of age (Fig 6L-M), 6 weeks after tamoxifen treatment is completed.

**Figure 6:**
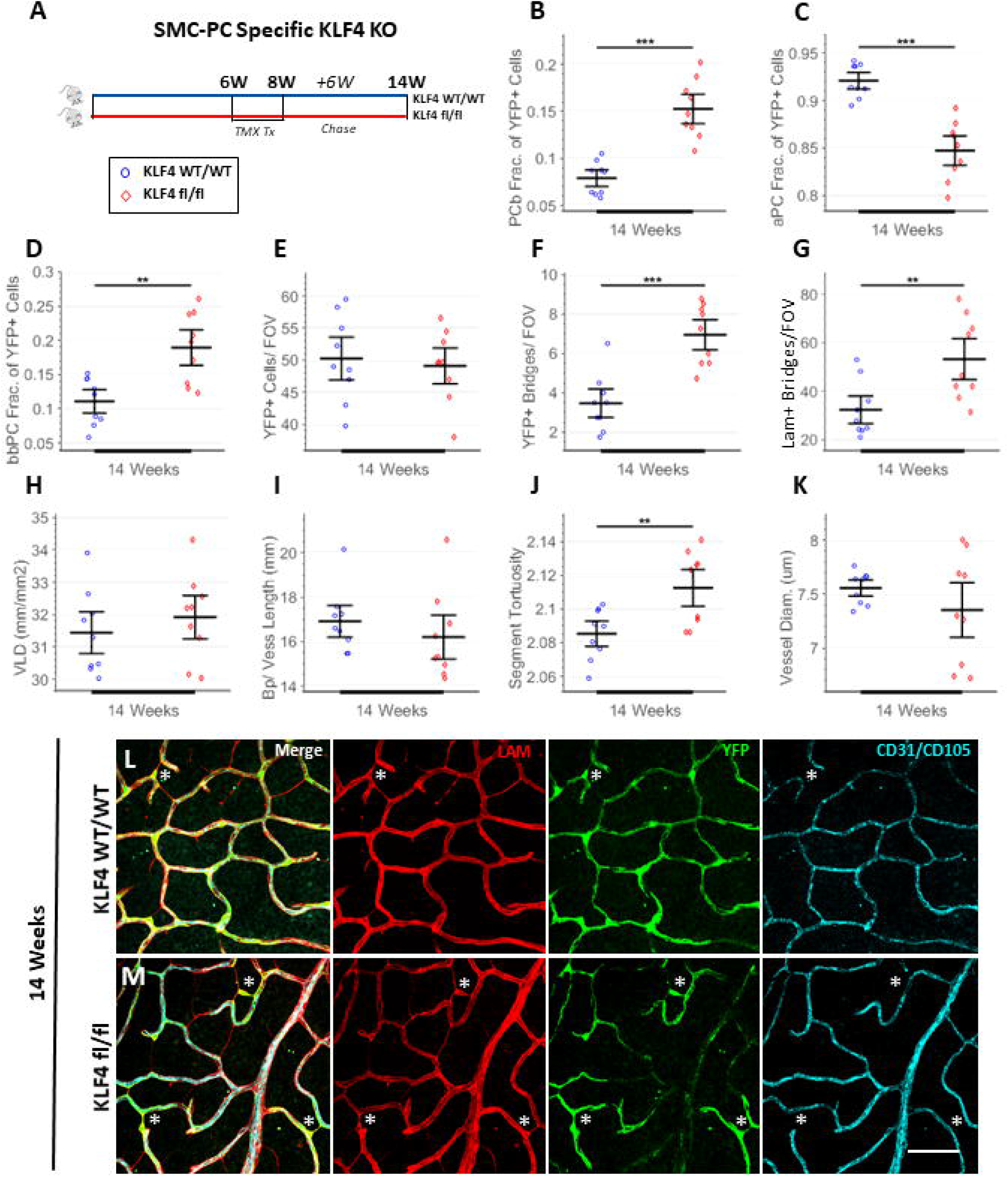
Loss of KLF4 in Myh11 lineage (Myh11-Lin(+)) cells exhibits enriched pericyte bridge density. (**A**) Experiment design. In the deep retinal plexus, quantification of Myh11-Lin(+) pericyte morphology, denoted by YFP expression, including (**B**) fraction of Myh11-Lin(+) cells with bridging phenotype, (**C**) fraction of My11-Lin(+) cells with attached phenotype, (**D**) fraction of My11-Lin(+) cells with basement membrane bridged phenotype, (**E**) total My11-Lin(+) cells per field of view, (**F**) My11-Lin(+) bridges per field of view, and (**G**) all laminin+ bridges per field of view. Vessel network morphology quantified with (**H**) vessel length density (mm/mm2), (**I**) number of segments per vessel length, (**J**) branch points per vessel length, and (**K**) vessel segment tortuosity (unpaired t-test, N=9 mice, 530 μm FOV). (**L, M**) Representative images of retinal deep plexus from each treatment group 6 weeks after tamoxifen induction, with anti-laminin (red), anti-YFP (green), anti-CD31 and anti-CD105 (cyan), with annotated pericyte bridges (star) (scale bar 50 μm).

### Pericytes are Capable of Migration and Process Extension Revealed Through In Vivo Time-lapse

While population level analysis demonstrated in the previous results suggest that pericyte bridges could be formed by active cell movement, there is a lack of evidence directly demonstrating that pericytes can migrate or extend off-vessel processes. As a requirement for pericyte remodeling, we investigate whether pericytes are capable of dynamic process extension and migration off-vessel, and explored this through *in vivo* time lapse imaging of corneal limbal vessels, a vascular bed bordering the sclera and cornea noted for its utility in live imaging^41^. To provide an angiogenic response mimicking that in diabetic retinopathy, silver nitrate burns were applied to the Myh11-RFP mouse cornea^41^, followed by pericyte tracking with the vascular perfusion of IB4 lectin to label vessels. At day 2 post-cornea burn, prior to when the majority of angiogenesis initiates in the model^42^, we observed multiple RFP+ cell somas migrating off the perfused vasculature (Fig. 7A-C) or RFP+ cell processes extending off-vessel (Fig 7D), confirming our hypothesis that pericytes are capable of undergoing migration and process extension dynamically in adult tissue.

**Figure 7:**
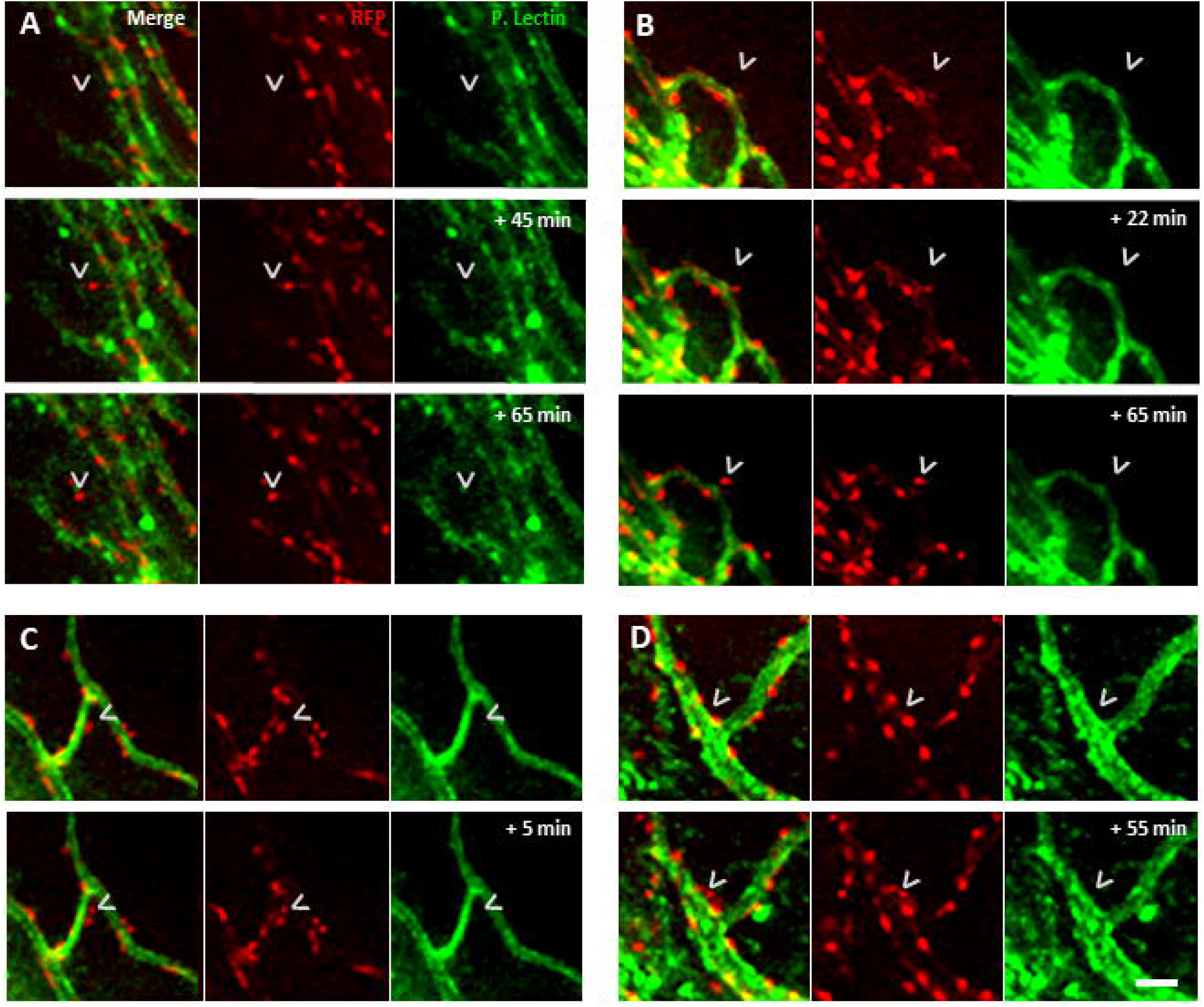
In limbal vessel network, Myh11-lineaged pericytes can detach processes and migrate off vessel visualized through in vivo time lapse. Timelapse imaging of corneal limbal vessels in Myh11-RFP mouse 2 days post cornea burn, with RFP+ Myh11-lineaged cells (red), and isolectin IB4 perfused vessels (green). (**A-C**) Time lapses that show an RFP+ cell soma starting fully associated with the vasculature and subsequently migrating off (arrow). (**D**) Timelapse of RFP+ cell extending a process off vessel (arrow) (scale bar 25 um).

## Discussion

The published literature has established the existence of pericyte bridge-like cells and basement membrane bridges in homeostasis and documented their elevated abundance in models of diabetes^10–13^. Here, we sought to determine if these cell and cell-free structures that span capillaries in the murine and human retina are reversibly modulated with stimuli relevant to diabetes and when a gene that is associated with cell migration^24, 40^ is ablated from pericytes. We show that: (1) pericyte-like bridging cells are, indeed, pericytes, and they retained their identity in the stimuli investigated, (2) the observed increase in pericyte bridges occurs in the absence of other microvascular remodeling events such as angiogenesis or regression that have been documented in hyperglycemia, (3) this increase in pericyte bridges is a reversible process by delivery of insulin, (4) pericytes are capable of active movement in adult vasculature and pericyte-specific deletion of KLF4, a transcription factor implicated in restricting cell migration, increases the abundance of pericyte bridges in the retina, and (5) pericyte bridges always colocalize with a basement membrane bridge, while endothelial cells never do.

Our conclusion that pericyte-like cells are indeed pericytes is established based on their traced lineage from Myh11 promoter activity and expression of Myh11 protein, a contractile myosin previously shown in vascularized tissues to exhibit exclusive expression in smooth muscle cells and pericytes^17^. Previous literature concluded that pericyte-like cells bridging capillaries were pericytes based on expression of NG2^14^ or PDGFRβ^18^, both nonexclusive cell markers. Our quantification of bridging cells used not only NG2 expression, but also colocalization with Col-IV or laminin basement membrane and lack of endothelial marker expression, suggests that they maintained their pericyte identity across cytokine and genetic perturbations. Across all stimuli, there were no NG2+ cell somas observed that lacked colocalization with Col-IV basement membrane, indicating that NG2 did not mark other cell types in these conditions.

Our data show that enrichment of pericyte bridges on the order of a few days in response to hyperglycemia occurs long before the vascular regression that has previously been observed at 6 months post STZ-induction^43^. This rapid change in pericyte bridges was recapitulated by injection of recombinant Ang2 and PDGF-BB, both of which are cytokines upregulated in diabetes^20,21^. Elevation of pericyte bridges in Akita mice^11^ provides further evidence that high blood sugar could be a short-term stimulus for enrichment of this pericyte phenotype.

For the first time, we show that the enrichment of pericyte bridges is a reversible phenomenon with restoration to basal levels following insulin treatment in STZ-induced hyperglycemia, both on a short-term time scale of days to a long-term timescale of months. While we assert that insulin exerts its primary effect through the normalization of blood glucose levels, we cannot rule out direct effects of insulin on pericytes or on endothelial cells in communication with pericytes. The dynamic capacity of pericyte bridges is reinforced by our observations that at long-term timepoints after exogenous delivery of Ang2 and PDGF-BB, pericyte bridges return to basal levels. Given that the management of blood sugar levels is known to have a strong influence on clinical outcomes in diabetic patients^44^, it is possible that the enrichment of pericyte bridges, as an early and reversible process, could serve as a target for early therapeutic interventions to slow or perhaps prevent diabetic vasculopathies.

Across all stimuli investigated, we showed there was no evidence of altered capillary density with appreciable effect size over the time-course of our studies, suggesting that the capillary network remains static during alterations in pericyte bridge density. Previous literature has hypothesized that pericyte bridges are formed either through the active movement of pericytes^10^, or as a result of vessel regression leaving behind an attached pericyte^9^. While our studies were not designed to rule out either of these hypotheses, our data is more consistent with pericyte movement being responsible for the pericyte bridge phenotype. Our time-lapse movie demonstrates that pericytes are capable of off-vessel migration and process extension over the timescale of minutes, far more rapid than pericyte movement demonstrated previously *in vivo*^45^. If vessels were regressing to create the enriched pericyte bridge density observed 4 days after injection of PDGF-BB and Ang2, an equal degree of angiogenesis would have to occur to maintain capillary density, yet without exception there were no signs of capillary sprouts or regression across all timepoints and stimuli. Our studies in STZ-induced diabetes also suggest that vessel regression is not implicated in the altered numbers of pericyte bridges because there were no changes in vascular density two weeks after STZ injection when pericyte bridge density was significantly elevated. Additionally, knockout of KLF4 in pericytes, a gene implicated in regulating cell migration^24,26^, resulted in enriched pericyte bridges with no appreciable change to pericyte or vessel density, further supporting that the occurrence of pericyte bridges does not require vessel regression. Additional experimentation is required, however, to definitively determine whether and to what extent vessel regression and/or pericyte migration causes the formation of pericyte bridges.

Both endothelial cells and pericytes are capable of producing basement membrane components such as laminin, fibronectin, and Col-IV^46^, and either cell type could be responsible for the formation of basement membrane bridges. If endothelial cells create them, it stands to reason that we would have observed an appreciable frequency of basement membrane bridges containing endothelial cells. However, across all stimuli, a subset of basement membrane bridges colabeled with pericytes but none with endothelial cells, and without exception all pericyte bridges colocalized with a basement membrane bridge. Enrichment of basement membrane bridge density in response to stimulus with PDGF-BB, a known pericyte chemokine that binds to PDGFRβ expressed in pericytes and not endothelial cells^39^, also implicates pericytes as the cells responsible for basement membrane bridges. The pericyte-specific knockout of KLF4 enriched basement bridges in tandem with pericyte bridges, further supporting the interpretation that basement membrane bridges are generated by pericytes which, in turn, could be mediated at least in part by pericyte migration.

Our results also reveal shortcomings in the classical retinal digest assay that has been cited as support for pericyte drop-out in diabetic conditions^20^, and they suggest that pericyte loss as a causative factor in diabetic vasculopathies needs to be reexamined, as it has already in the Akita mouse model^11^. Our analysis of immunostained retinas revealed that up to 50% of all pericyte somas were associated with a basement membrane bridge (combination of pericyte bridges and basement membrane bridged pericytes), which would have been miscounted as endothelial somas^47,48^ in the retinal digest assay and potentially accounts for the approximately 30%^20^ loss in pericyte density observed with this assay in diabetes and Ang2 stimulation. We show that pericytes only occupy a subset of basement membrane bridges, but this cell-specific colocalization is not captured with the histological staining used in retinal digests, limiting the assay’s usefulness for quantifying pericyte bridges and total pericyte cell count.

The existence of pericyte bridges in homeostatic conditions that are known to gradually decrease with age^9^ suggests that this pericyte subpopulation serves a homeostatic function. The enrichment of pericyte bridges during diabetic conditions could alter pericyte coverage of the network^49^ and suggests that they could play a contributing or protective role in vasculopathy. Elevated pericyte bridges may prime the vessel network for remodeling, and basement membrane bridges are hypothesized to offer a preferential route for the rapid growth of new blood vessels^50^. Earlier events in disease progression are thought to be superior avenues for interventions since preventative treatments avoid the need to regenerate damaged tissue. Modulation of pericyte bridges in hyperglycemia could provide a novel cell behavior as a potential therapeutic target.

## Author Contributions

B.A.C designed and performed experiments and drafted figures and manuscript with input from all authors. R.D. developed software for data analysis. C.M. aided with data analysis, acquisition, and design of experiments. H.C.R. and N.S. aided with design of experiments and acquisition of data. K.F., R.P. aided with data analysis and acquisition. M.K.G. aided with development of project and in vivo imaging. J.C, L.M. and G.K.O aided with developing central thesis of manuscript. P.Y., and S.M.P. supervised the project. All authors discussed the results and contributed to the final manuscript.

## Acknowledgements

We would like to thank vivarium staff for helping to maintain the mouse strains used for this research, along with Anthony Bruce managing Dr. Peirce-Cottler’s lab. We are also grateful for the Advance Microscopy Facility at UVA for providing the equipment necessary for high resolution confocal imaging, and Hamzah Shariff and Brian Rothemich for their help with data analysis. B.A.C and S.M.P are guarantors for this manuscript.

## Funding

NIH R21 EY028868-01, NIH-U01AR069393, NIH-U01HL127654, The Hartwell Foundation, The Stanford Allen Discovery Center (to S.M.P.).

## Data and Resource Availability

The datasets generated during and/or analyzed during the current study are available from the corresponding author upon reasonable request. The mouse strains generated during and/or analyzed during the current study are either commercially available or available through the labs that generated them.

## Conflict of Interests

P.A.Y.: RetiVue, LLC (Personal Financial Interest/Employment), Genentech/Roche (Consultant).

