## Supplement File for "Pericyte Bridges in Homeostasis and Hyperglycemia: Reconsidering Pericyte Dropout and Microvascular Structures"

**Short Running Lead**: Pericyte Bridges in Diabetes and Homeostasis

**Corresponding Author**:

Shayn M. Peirce, Ph.D.

Professor

Department of Biomedical Engineering

415 Lane Road

University of Virginia

Charlottesville, VA 22908

**Supplement Contains**

Supplementary Figures 1-7

Supplementary Data 1-3

Supplementary Materials 1-5

Supplementary Tables 1-2

Supplementary Timelapse Movies 1-2

### Supplementary Figures

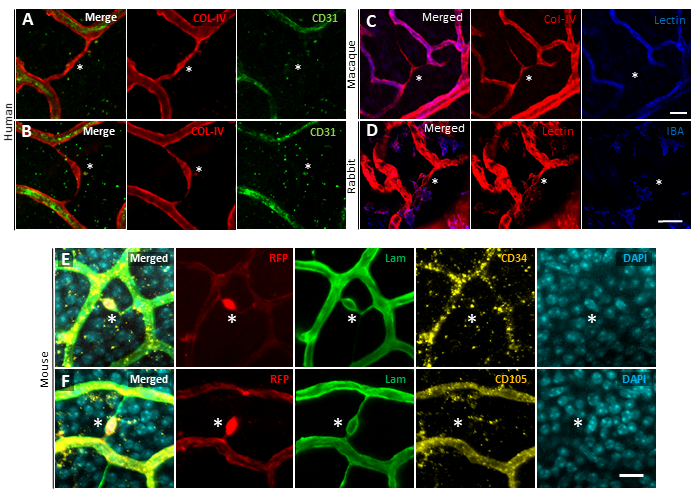

**Supplementary Figure 1: Off-vessel Col-IV tracks are found in various mammals, and do not express the endothelial markers CD34 and CD105 in mice**. (**A, B**) Col-IV track in human tissue visualized with anti-Col-IV (red) and anti-CD31 (green). (**C**) Col-IV track visualized in maquake retina with anti-Col-IV (red) and isolectin IB4 (blue). (**D**) Col-IV track visualized in rabbit retina with anti-Col-IV (red) and anti-IBA1 (blue) (scale bar 25 um). (**E**) Murine retina of Myh11-RFP mice, with RFP denoting Myh11 lineage-marked cells (red, star), anti-laminin (green), anti-CD34 (yellow), and DAPI nuclei (cyan). (**F**) Murine retina with RFP denoting lineage-marked cells (red, star), anti-laminin (green), anti-CD105 (yellow), and DAPI nuclei (cyan) (scale bar 15 um).

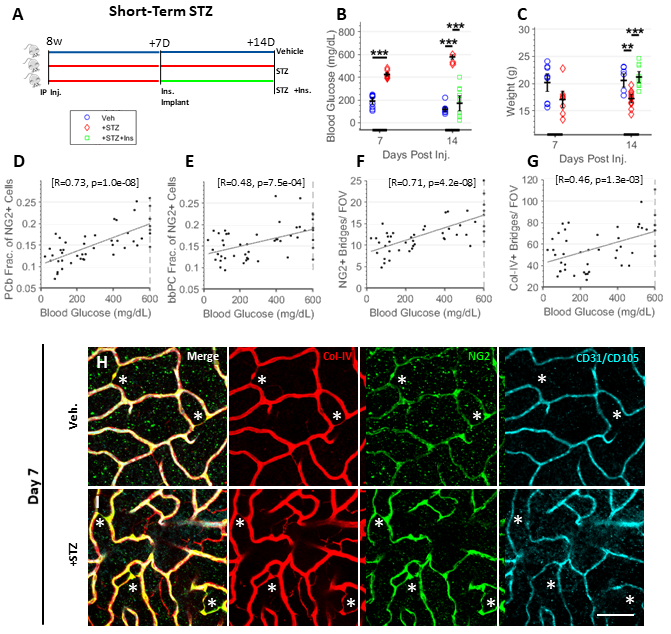
**Supplementary Figure 2:**

**Density of pericyte bridges and basement membrane bridges correlate with blood glucose level at time of sacrifice with short-term diabetes.** (**A**) Experiment design illustrating study groups and timepoints. (**B**) Blood glucose concentrations and (**C**) weight at time of sacrifice for each treatment group (day 7: unpaired t-test, day 14: 1-way ANOVA with Tukey multiple comparisons, N=9 mice). For all mice across treatment groups, Pearson correlation between blood glucose level at harvest and (**D**) fraction of NG2+ pericyte bridges, (**E**) fraction of NG2+ pericytes with basement bridged phenotype, (**F**) NG2+ basement membrane bridges per field of view, and (**G**) all Col-IV+ bridges per field of view (Pearson R value and p value in brackets). (**H, I**) Representative images of retinal deep plexus at day 7 post STZ injection from each treatment group, with anti-Col-IV (red), anti-NG2 (green), along with anti-CD31 and anti-CD105 (cyan), with annotated IBCs (star) (scale bar 50 um).

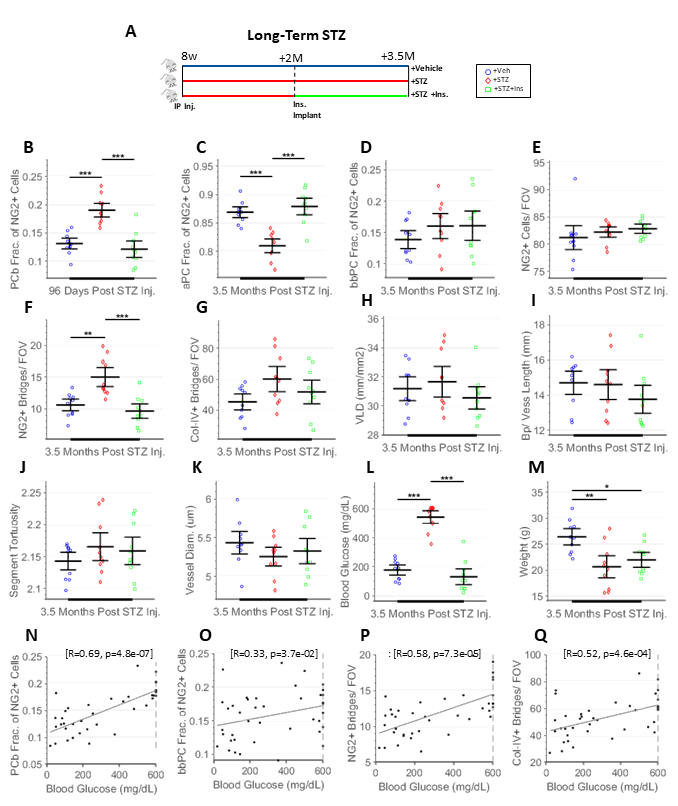

Supplementary Figure 3: **Long-term STZ-induced hyperglycemia has enriched pericyte bridge density normalized with insulin treatment over a static vessel network structure**. (**A**) Experiment design. In the deep retinal plexus, quantification of pericyte morphology, including (**B**) fraction of NG2+ cells with pericyte bridge phenotype, (**C**) fraction of NG2+ cells with attached pericyte phenotype, (**D**) fraction of NG2+ cells with basement bridged phenotype, (**E**) total NG2+ cells per field of view, (**F**) NG2+ bridges per field of view, and (**G**) all Col-IV+ bridges per field of view. Vessel network morphology quantified with (**H**) vessel length density (mm/mm2), (**I**) branch points per vessel length, (**J**) vessel segment tortuosity, and (**K**) vessel diameter (1-way ANOVA, Tukey multiple comparisons, N=10 mice, FOV 530 µm). (**L**) Blood glucose concentrations and (**M**) weight at time of sacrifice for each treatment group (1-way ANOVA, Tukey multiple comparisons, N=10 mice). For all mice across treatment groups, Pearson correlation between blood glucose level at harvest and (**N**) fraction of NG2+ pericyte bridges, (**O**) fraction of NG2+ pericytes with basement bridged phenotype, (**P**) NG2+ basement membrane bridges per field of view, and (**Q**) all Col-IV+ bridges per field of view (Pearson R value and p value in brackets).

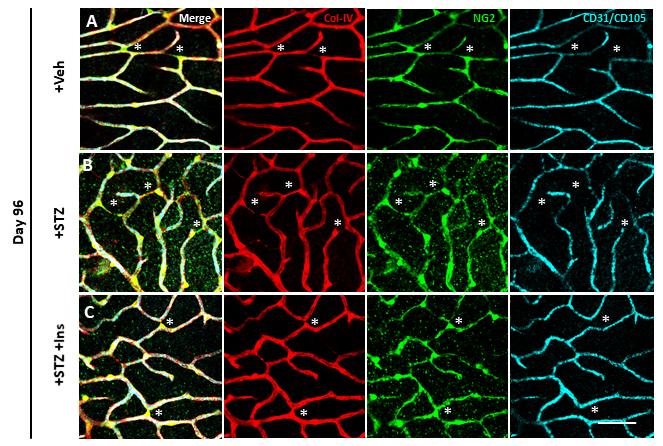

**Supplementary Figure 4:** **Representative images of vasculatures subject to long-term hyperglycemia**. (**A-C**) Representative images of retinal deep plexus at 3.5 months from each treatment group, with anti-COL-IV (red), anti-NG2 (green), along with anti-CD31 and anti-CD105 (cyan), with annotated pericyte bridges (star) (scale bar 50 um).

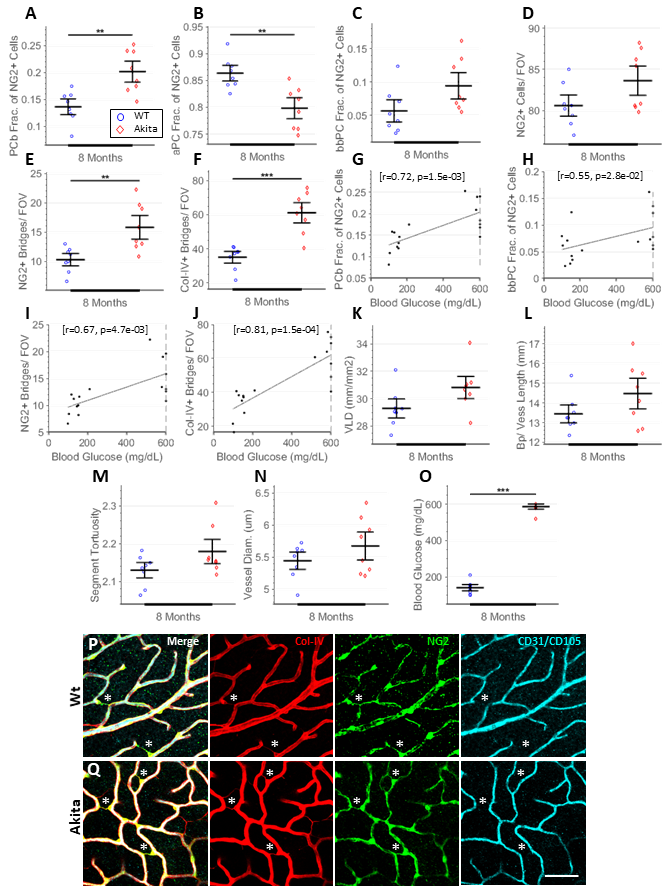

**Supplementary Figure 5:** **Akita mice over the long-term (8 months) exhibit enriched density of pericyte bridges over a morphologically stable vessel network structure with minor remodeling**. Quantification of pericyte morphology, including (**A**) fraction of NG2+ cells with pericyte bridge phenotype, (**B**) fraction of NG2+ cells with attached phenotype, (**C**) fraction of NG2+ cells with basement bridged phenotype, (**D**) total NG2+ cells per field of view, (**E**) NG2+ off vessel basement membrane bridges per field of view, and (**F**) all Col-IV+ bridges per field of view (unpaired t-test, N=8, FOV 530 µm). For all mice across study groups, Pearson correlation between blood glucose level at harvest and (**G**) fraction of NG2+ cells with pericyte bridge phenotype, (**H**) fraction of NG2+ cells with basement membrane bridged phenotype, (**I**) NG2+ off-vessel bridges per field of view, and (**J**) all Col-IV+ bridges per field of view (Pearson R value and p value in brackets). Vessel network morphology quantified with (**K**) vessel length density (mm/mm2), (**L**) branch points per vessel length, (**M**) vessel segment tortuosity, and (**N**) vessel diameter. (**O**) Blood glucose level at time of sacrifice from both groups to confirm diabetic phenotype. (**P, Q**) Representative images of retinal deep plexus from each treatment group at 8 months of age, with anti-Col-IV (red), anti-NG2 (green), anti-CD31 and anti-CD105 (cyan), with annotated pericyte bridges (star) (scale bar 50 um).

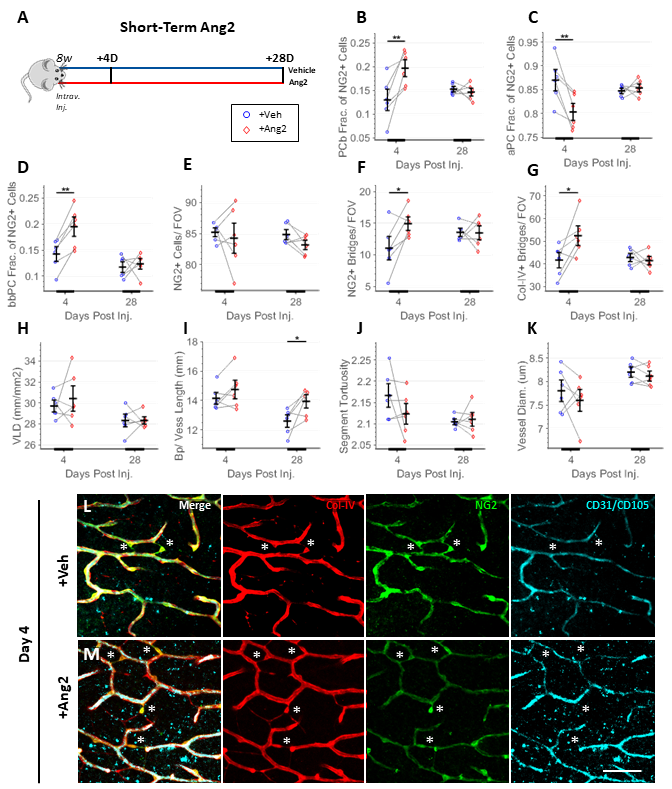

**Supplementary Figure 6: Intravitreal injection of Ang2 exhibits transiently enriched pericyte bridge density over a morphologically static vessel network**. (**A**) Experiment design. In the deep retinal plexus, quantification of pericyte morphology, including (**B**) fraction of NG2+ cells with pericyte bridge phenotype, (**C**) fraction of NG2+ cells with attached pericyte phenotype, (**D**) fraction of NG2+ cells with basement bridged phenotype, (**E**) total NG2+ cells per field of view, (**F**) NG2+ off vessel bridges per field of view, and (**G**) all Col-IV+ bridges per field of view (N=6 mice). Vessel network morphology quantified with (**H**) vessel length density (mm/mm2), (**I)** branch points per vessel length, (**J**) vessel segment tortuosity, and (**K**) vessel diameter (paired t-test at each timepoint, N=6 mice, 530 µm FOV). (**L, M**) Representative images of retinal deep plexus at day 4 from each treatment group, with anti-Col-IV (red), anti-NG2 (green), along with anti-CD31 and anti-CD105 (cyan), with annotated pericyte bridges (star) (scale bar 50 um).

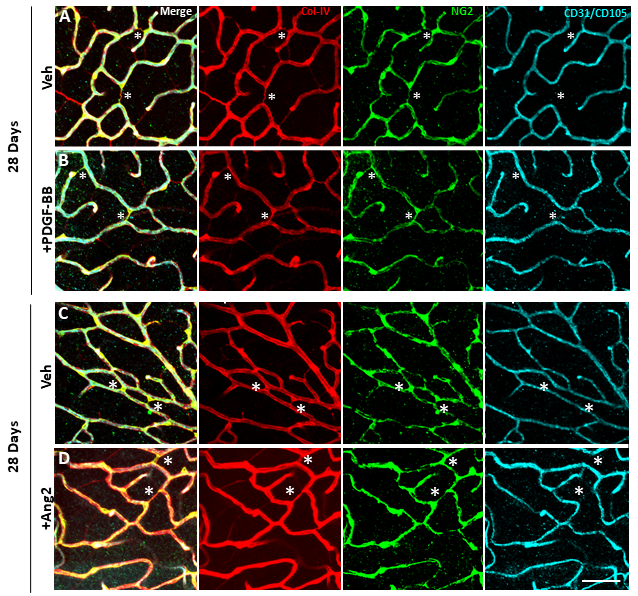

**Supplementary Figure 7: Vascular network is normalized 28 days post injection of PDGF-BB and Ang2.** (**A, B**) representative images from both study groups 28 days after injection of PDGF-BB, with anti-Col-IV (red), anti-NG2 (green), along with anti-CD105 and anti-CD31 (cyan). (**C, D**) representative images from both study groups 28 days after injection of Ang2, with anti-Col-IV (red), anti-NG2 (green), along with anti-CD105 and anti-CD31 (cyan), with pericyte bridges annotated (star) (scale bar 25 µm).

### Supplementary Data

#### Supplementary Data 1: Long-Term Hyperglycemia Elevates while Insulin Treatment Reduces Pericyte Bridge Density in STZ-induced Diabetes

As with short-term diabetes, perivascular remodeling was observed over the long-term at 3.5 months post STZ injection, and insulin treatment after the second month resulted in partial rescue of diabetic perivascular phenotype (Supplementary Fig. 3A-C), suggesting that these phenotypes are not merely associated with the short-term initiation of hyperglycemia. The density of pericyte bridges with diabetic mice was enriched by 31.0% (Supplementary Fig. 3B), while attached pericyte density was reduced 7.3% (Supplementary Fig. 3C, p=2.97E-5). Insulin treatment returned densities to normal with pericyte bridges reduced 57.0% and attached pericytes increased 7.9% (p=2.63E-6). There was an insignificant elevation in basement bridged pericytes in diabetes and insulin treatment (Supplementary Fig. 3D, p=0.368). No change was observed in total density of all NG2+ pericytes (Supplementary Fig. 3E, p=0.465). Basement bridges labeled with Col-IV and the NG2 co-labeled subset displayed similar trends as those found with pericyte bridge density across study groups (Supplementary Fig. 3F-G). There was no evidence of angiogenesis or regression, as evidenced by no change to vessel length density (Supplementary Fig. 3H, p=0.391), branchpoints per vessel length (Supplementary Fig. 3I, p=0.344), segment tortuosity (Supplementary Fig. 3J, p=0.420), or vessel diameter (Supplementary Fig. 3K, p=0.389). In support of perivascular remodeling, the density of pericyte bridges, basement bridged pericytes, and off vessel bridges all correlated with blood glucose of each mouse at time of sacrifice (Supplementary Fig. 3N-Q). Blood glucose and mouse weight confirms hyperglycemia for each study group (Supplementary Fig. 3L-M); representative images at 3.5 months provided (Supplementary Fig. 4A-C).

#### Supplementary Data 2: Genetic Knockout of Insulin in Akita Mouse Elevates Pericyte Bridge Density

In the Ins2Akita genetic mouse model, the fraction of NG2+ cells with pericyte bridge phenotype was enriched 32.4% at 8 months compared to genetic background (WT) in tandem with an 8.2% decrease in the fraction of NG2+ on-vessel attached pericytes (Supplementary Fig. 5A-B, p=1.97E-3). There was a 39.9% increasing trend in the fraction of NG2+ cells with basement bridge pericyte phenotype (Supplementary Fig. 5C, p=0.0595). Total NG2+ cells had an enriched trend with a small effect size of 3.6% (Supplementary Figure 5D, p=0.0720). There was a 34.8% increase in NG2+ basement membrane bridges (Supplementary Fig. 5E, p=4.22E-3) and a 42.6% increase in all Col-IV+ bridges (supplementary Fig 5F, p=9.89E-5). There was a significant correlation between blood glucose at time of sacrifice and density of pericyte bridges, basement bridged pericytes, and basement membrane bridges (Supplementary Fig. 5G-J, Pearson correlation). While there was a minor 4.9% enriched trend in vessel length density between Akita and WT (Supplementary Fig. 5K, p=0.0627), all other metrics characterizing the vasculature showed no change, including branchpoints normalized to vessel length (Supplementary Fig. 5L, p=0.128), segment tortuosity (Supplementary Fig. 5M, p=0.866), and vessel diameter (Supplementary Fig. 5N, p=0.224). Blood glucose confirmed the diabetic phenotype (Supplementary Fig. 5O) seen in representative images (Supplementary Fig. 5P-Q) provided.

#### Supplementary Data 3: Injection of Recombinant Ang2 Transiently Elevates Pericyte Bridge Density

Four days post-Ang2 injection (Supplementary Fig. 6A), pericyte bridge density was transiently enriched 33.9% relative to vehicle control (Supplementary Fig. 6B) and attached pericytes were reduced 8.3% (Supplementary Fig. 6C, p=9.72E-3). At day 28, both pericyte subpopulations recovered to basal levels (p=0.571). Pericytes with a basement bridged phenotype exhibited the same pattern, with 26.9% enrichment at day 4 (Supplementary Fig. 6D, p=1.88E-3) and restoration at day 28 (p=0.697). Total NG2-labeled cell density remained constant both at days 4 (Supplementary Fig. 6E, p=0.647) and 28 (p=0.957). Basement membrane bridges labeled with Col-IV and the subset colabeled with NG2 followed similar trends as pericyte bridge density across study groups (Supplementary Fig. 6F-G). No angiogenesis was observed at day 4, with vessel length density (Supplementary Fig. 6H, p=0.525), branchpoints per vessel length (Supplementary Fig. 6I, p=0.158), segment tortuosity (Supplementary Fig. 6J, p=0.225), vessel diameter (Supplementary Fig. 6K, p=0.423) remaining unchanged. Representative images from day 4 (Supplementary Fig. 6L, M) and day 28 are provided (Supplementary Fig. 7C-D).

### Supplementary Materials

#### Supplementary Materials 1: Mouse Treatments

With the lineage tracing mice used, cre recombinase was activated in male mice with a series of ten 1-mg intraperitoneal injections of tamoxifen (Sigma, cat. #. T-5648,) from 6 to 8 weeks of age, for a total of 10 mg tamoxifen/mouse. Animals were allowed to recover for ≥4 weeks to allow residual tamoxifen to leave the system before experiments/tissue isolation.

For STZ induced diabetes, at 7 weeks of age, glucose was taken for all mice after a 2 hour fast to establish a baseline glucose level. At 8 weeks, mice were injected interperitoneally with filter sterilized 200 mg/kg STZ (Sigma, Cat. # 85882) in 0.1 M Citrate buffer to initiate severe hyperglycemia, or simply given vehicle control. Mice were fasted for 4 hours prior to injection, and provided with 10% sucrose for drinking water overnight. Glucose was again taken a week after injection after a 2 hour fast to confirm hyperglycemia with blood glucose above 300 mg/dL. For the duration of experiment, blood glucose and body weight were checked twice a week with a glucose meter (Bayer, Pittsburgh, PA, cat. # 9628), and diabetic mice were injected with up to 1 unit of Humulin-N (Ely Lilly, USA) as necessary to maintain body weight. Maximum measurement for glucose meter saturated at 600 mg/dL, so measurements about that value were recorded as max value. To examine the effects of normalizing glucose levels in diabetic mice and eliminating the effects of vascular injury from STZ directly causing vascular remodeling, the insulin treatment group was implanted with Linbit insulin implants (Linbit, Cat. # Pr-1-B, Ontario, Canada) via subcutaneous injection following manufacturer’s instructions. Additional implants were added through time to bring blood glucose levels below 300 mg/dL in two consecutive readings in order to normalize hyperglycemia. Mice without insulin implants (STZ diabetic and vehicle) were given sham injections to mimic the injury from implantation.

#### Supplementary Materials 2: Intravitreal Injections of Ang2 and PDGF-BB

Mice were anesthetized with using inhalation of 2% isoflurane/oxygen (Henry Schein Animal Health, Dublin, Ohio, cat. # 029405). A drop of sterile 0.5% Proparacaine hydrochloride ophthalmic solution (Henry Schein Animal Health, Dublin, Ohio, cat. # 1127199) was added as a topical anesthetic to numb the eye before injection. For the Ang2 experiments, mice were intravitreally injected in one eye with 1.5 µL of 0.67 µg/µL of carrier-free recombinant human Ang2 (R and D Biosystems, Minneapolis, MN, cat. # 623-AN-025/CF), in PBS, and the other eye 1.5 µL of PBS to act as contralateral control. For the PDGF-BB experiments, mice were intravitreally injected in one eye with 1.5 µL of 2 µg/µL of carrier free recombinant mouse PDGF-BB (Shenandoah Biotechnology, Warwick, PA, cat. # 200-58-100ug) in sterile water, and the other eye 1.5 µL of sterile water to act as contralateral control. To carry out intravitreal injections, a 30G needle (Becton Dickerson, Franklin Lakes, NJ, cat. # 305128) was used to initially poke through the sclera, followed the injection with a Hamilton syringe (Hamilton Company, Reno, NV 7633-01,65 RN) with a Hamilton 33G needle (Hamilton Company, Reno, NV, cat. #7803-05, PT4).

#### Supplementary Materials 3: Retina Harvest and Immunostaining

Mice were sacrificed via CO2 asphyxiation with cervical dislocation for secondary sacrifice, eyes enucleated, and incubated in 4% PFA for 10 minutes, and retinas surgically isolated. Tissue was permeabilized with 1 mg/mL digitonin for 2 hours (Sigma, cat. # D141-500MG), immunostained with the same concentration of digitonin for all steps left overnight at 4 °C, and washed with 0.1% saponin solution in between (Sigma, cat. # 84510-100G). Primary antibodies are listed in Supplementary Table 1, and secondary antibodies in Supplementary Table 2. For staining with NG2 and Col-IV, retinas were subject to proteinase-K digestion after fixation to aid with antibody penetration (see main text for reference).

#### Supplementary Materials 4: Quantifying Microvascular Structure and Pericyte Phenotype

Cell counts of pericyte association state with the vasculature was quantified using Fiji’s Cell Counter plugin in a blinded fashion, along with a blind rating of image and staining quality to unbiasedly filter out images with poor image quality. Vessel structure was analyzed with software written in MATLAB implementing algorithms and metrics developed previously to quantify images of vascular networks. In brief, the images where thresholded by subtracting a heavily blurred image that captured the background features from the original image to acquire a background subtracted image. Pixel intensities were thresholded based on a global threshold value, and a series of smoothing filters used on the segmented image to smooth the border of the segmentation, fill in holes in connected components, and remove spurious segmented pixels. The user is presented with this segmentation and allowed to make edits with the image: either removing segmented structures that do not capture the vasculature, or adding structures where the segmentation failed to segment the vasculature. While performing curation on the image, the image name is obfuscated to blind the user and keep processing unbiased. The vessel centerline is calculated with skeletonization, and spurious skeleton segments are removed, and branchpoints and endpoints identified. Several metrics were used to quantify vessel network morphology as done previously^68^. Vessel area fraction is the fraction of pixels of the foreground vasculature out of total image area. Vessel length density is the length of the vessel centerline, converted to millimeters using the resolution of the image, divided by the area of the image. Vessel diameter is calculated by taking the Euclidean distance transform of the complemented vessel segmentation, and selecting pixels that overlap the vessel centerline, and then doubling those values and adding one. Branchpoints per vessel length is the number of branchpoints in the image divided by length of centerline in millimeter units. Tortuosity was measured based on average of vessel segment length normalized to distance between vessel segment endpoints.

Pericytes were defined by NG2 cell soma that colocalized with a Col-IV basement membrane- for all stimuli investigated, there were no NG2+ cell somas that were negative for Col-IV. Pericytes were split into several subpopulations based on their morphology relative to the vasculature. Pericytes that had all cell processes and cell soma attached to a vasculature was classified as an attached pericyte. This subpopulation was split further between pericytes connected to an empty basement membrane bridge (negative for endothelial marker and pericyte marker) and those that were not. Pericytes that had a process or cell soma bridging capillaries were classified as pericyte bridges. For pericyte subpopulation densities, the fraction of pericyte bridges is reported as well as the fraction of attached pericytes. Note that this is redundant data (same p values for both subpopulations) since they are fractions that together form the total pericyte population. We reported both values to convey the relative change in effect sizes across study groups.

#### Supplementary Materials 5: Data Acquisition, Sampling, and Statistics

For the investigation of basement membrane bridges in homeostatic retina, an unpaired two-tailed welches’ t-test was used for bridge marker expression and 1-way Kruskal-Wallis with Bonferroni multiple comparison (N=6 mice). For the short-term diabetes experiment, a two-tailed unpaired t-test was used to evaluate cell count and vessel architecture morphologies at day 7, while a 1-way ANOVA with Tukey’s multiple comparisons test was used for day 14 (N=8). For the long term STZ experiment, a 1-way ANOVA was used with Tukey’s multiple comparisons (N=10). For the Ang2 (N=6) and PDGF-BB (N=10) intravitreal injection experiment, a paired t-test was used at day 4 and day 14. For the KLF4 knockout experiment, an unpaired 2-tailed t-test was used (N=9). For the Akita experiment, an unpaired 2-tailed t=test was used (N=8). All images were acquired from deep retinal plexus approximately mid-distance radially between the optic disk and peripheral edge of the retina. For experiments with contralateral eye control, 4 fields of view per retina, 530 µm square, were acquired of the retina deep plexus, at an intermediate radial distance between the optic disk and peripheral retina. For experiments without a contralateral control, 5 fields of view per mouse were used for quantification.

For live imaging, animals were anesthetized with an intraperitoneal injection of ketamine/xylazine/atropine (60/4/0.2 mg/kg body weight) (Zoetis; Kalamazoo, MI/West-Ward; Eatontown, NJ/Lloyd Laboratories; Shenandoah, IA). A drop of sterile 0.5% Proparacaine hydrochloride ophthalmic solution was added as a topical anesthetic to numb the eye before injection. To allow visualization of vascular endothelium, anesthetized mice were administered a retro-orbital injection of labeled isolectin (IB4-Alexa647; Life Technologies, Carlsbad, CA) 30 minutes before sacrifice. Animals were then mounted on a microscope stage and imaged with a Nikon point scanning confocal (Nikon Instruments Incorporated, Melville, NY; Model TE200-E2; 10X air objective) while kept unconscious with isoflurane anesthesia and a heating pad to maintain body weight. Mice were sacrificed at the end of the imaging session. For histological imaging, high resolution imaging was carried out on a Zeiss LSM 880 with a 63x Plan-Achromat/1.40 oil DIC. Imaging for quantification of vasculature and pericyte phenotype was carried out with a Nikon point scanning confocal (Nikon Instruments Incorporated, Melville, NY; Model TE200-E2) with a Plan Flour 20X/0.75 oil immersion objective.

### Supplementary Tables

#### Table 1: Primary Antibodies

| **Target/Name** | **Imm. Species** | **Host Species** | **Company** | **Cat #** | **Dilution** |
| --- | --- | --- | --- | --- | --- |
| Anti-NG2 | Rat | Rabbit | Millpore, Darmstadt, Germany | AB5320 | 1:200 |
| Anti-Collagen-IV | Human | Goat | Bio-rad, Oxford, UK | 13400 | 1:100 |
| Isolectin GS-IB4 | - | Griffonia simplicifolia | Life Technologies, Carlsbad, CA | I32450 | 1:150 |
| DAPI | - | - | ThermoFisher Scientific, Waltham, MA | D1306 | 1:500 |
| Anti-CD34 | Mouse | Rat | Biolegend, San Diego, CA | 119301 | 1:200 |
| Anti-CD105 | Mouse | Rat | Biolegend, San Diego, CA | 120402 | 1:200 |
| Anti-Laminan [AF488] | Mouse | Rabbit | Novus, Saint Charles, MO | NB300-144AF488 | 1:100 |
| Anti-Fibronectin | Mouse | Rabbit | Abcam, Cambridge, UK | AB2033 | 1:200 |
| Anti-RFP | - | Rabbit | Abcam, Cambridge, UK | AB62341 | 1:200 |
| Anti-CD31 | Human | Mouse | Biolegend, San Diego, CA | 910005 | 1:200 |
| Anti-CD31 | Mouse | Rat | Biolegend, San Diego, CA | 102504 | 1:300 |
| Anti-Myh11 | Mouse | Rat | Kamiya Biomedical Company, Seattle, WA | MC-352 | 1:300 |
| Anti-GFP [AF647] | *Aequorea victoria* | Rabbit | Life Technologies, Carlsbad, CA | A-31852 | 1:150 |
| Anti-IBA1 | Rat | Goat | Abcam, Cambridge, UK | AB107159 | 1:200 |

#### Table 2: Secondary Antibodies

| **Host** | **Target** | **Flour** | **Company** | **Cat #** | **Dilution** |
| --- | --- | --- | --- | --- | --- |
| Donkey | Rabbit IgG | AF546 | Invitrogen, Carlsbad, CA | A10040 | 1:600 |
| Donkey | Rabbit IgG | AF647 | ThermoFisher, Waltham, MA | A-31573 | 1:600 |
| Donkey | Rat IgG | AF488 | Invitrogen, Carlsbad, CA | A-21208 | 1:600 |
| Donkey | Rat IgG | AF647 | Abcam, Cambridge, United Kingdom | AB150155 | 1:600 |
| Donkey | Goat IgG | AF546 | Invitrogen, Carlsbad, CA | A11056 | 1:600 |
| Donkey | Goat IgG | AF647 | Invitrogen, Carlsbad, CA | A21447 | 1:600 |

### Online Supplemental Videos 1-2

Included in online supplementary data are files “Online Supplemental Video 1.mp4” and “Online Supplemental Video 2.mp4”, both of which are in vivo timelapse movies of the corneal limbal vessel network 2 days post silver nitrate cornea burn. Images were acquired on a Nikon scanning CLSM with a 10X/0.3 objective. Pericytes were endogenously labeled with tdTomato in the Myh11-RFP mouse model, and vessels were labeled with perfused IB4 lectin. Each frame of the videos are approximately 2.5 minutes apart in time.
